# Mutation accumulation underpins evolution of lifespan extension by dietary restriction

**DOI:** 10.1101/2025.06.18.660314

**Authors:** Sara D. Irish, Annabel Kimberley, Simone Immler, Jacob Moorad, Alexei A. Maklakov

## Abstract

Dietary restriction (DR) extends lifespan in animals and plants, but its evolutionary causes are elusive. Adaptive hypotheses posit that DR extends lifespan because organisms reallocate resources from reproduction to survival (‘disposable soma’) or recycle cellular waste to maximize either their immediate reproduction (‘nutrient recycling’) or survival (‘clean cupboards’). We developed an experimental paradigm that tricks *Caenorhabditis elegans* nematodes into increasing their reproductive effort under DR via food odour, thus allowing us to test these hypotheses. We found that experimentally increased reproduction under DR does not affect immediate or long-term survival benefits compared to DR animals that did not reproduce, thus refuting all three adaptive hypotheses. Our data suggest that a large part of suppressed fertility under DR is a result of organisms refraining from producing offspring in a poor environment. We developed a model based on Hamiltonian forces of selection to show that lifespan extension under DR evolves because DR suppresses fertility, directly increasing selection against mortality in DR environment. Our analytical approach suggests that DR-driven lifespan extension can evolve under a broader range of conditions not previously anticipated, such as a relaxed need for physiological or genetic trade-offs. Instead, we show how reduced survival on plentiful food can evolve via mutation accumulation.

## Introduction

Dietary restriction (DR), the reduction of nutrients without malnutrition, is well known to extend lifespan across a broad range of taxa, from plants to primates^1–4^, yet the evolutionary causes of this phenomenon remain poorly understood. While it has been well established that DR extends lifespan, this is not without cost to individuals, as limiting nutrient intake also induces declines in fertility. This has led to much debate around the mechanisms through which DR works and whether the response to DR carries adaptive benefits that would translate to natural populations.

DR may extend lifespan when organisms make an adaptive ‘decision’ to delay reproduction when nutrients are scarce and to allocate remaining resources to somatic maintenance until food becomes available again (the ‘resource reallocation’ theory) ^5,6^. While this theory enjoyed some support, many empirical studies in recent decades showing that increased lifespan and reduced reproduction under DR can be uncoupled.

Alternatively, the ‘nutrient recycling’ hypothesis suggests that lifespan extension via DR occurs through the recycling of extraneous proteins and cells, but for the purpose of maximising immediate reproduction rather than for survival. This views increased longevity as an unselected by-product of increased recycling ^7^. However, it is argued that delaying reproduction is an ineffective strategy for maximising fitness due to higher rates of extrinsic mortality in nature, especially in short-lived organisms ^8,9^. Finally, the ‘clean cupboards’ hypothesis argues that when nutrients are scarce, individuals will use up damaged cells and misfolded proteins to make up for the deficiency to maximise immediate survival, while improved late-life survival is again seen as unselected by-product of the removal of metabolic waste^10^. While some evidence supports both of these models^11^, other studies find that DR animals not only live longer than controls^12^ but also exhibit improved health and increased tolerance to abiotic stress and pathogens^2,13^. This suggests that improved somatic maintenance under DR response may be beneficial for fitness under a broad range of ecologically relevant conditions.

A more parsimonious explanation for the evolution of DR is that different environments manifest different selection gradients for vital rates. Specifically, the direct effect of DR is to suppress reproduction, and this has an indirect effect of increasing selection gradients for survival, especially at late age. Provided that some genes are expressed differently in different environments, standard evolutionary theory predicts that selection will result in the evolution of longer lifespan in the environment that suppresses reproduction. The evolution of lifespan extension under DR does not require resource allocation trade-offs but rather stems directly from the concept of ‘selection shadow’, the declining force of natural selection with advancing age^14,15^, coupled with differential gene expression. Because adult survival is more important for evolutionary fitness in DR environment compared to non-DR environment, the DR response – the difference in somatic maintenance between DR and non-DR environments – can evolve via accumulation of germline mutations that reduce somatic maintenance in non-DR environments.

Here we tested previous models for the evolution of the DR response in *Caenorhabditis elegans* nematodes using an experimental paradigm that allows us to induce reproduction in DR conditions. Environmental sensing plays an important role in the DR response^16–18^, and olfactory and gustatory neurons act through partially independent pathways to influence lifespan extension^16,17^. Previous work has shown that in *C. elegans,* introduction of the scent of food in DR conditions can partially restore reproduction at the cost of lifespan thus supporting resource reallocation theory^19^. However, we developed an experimental protocol where we can substantially increase reproduction under DR without any effect on lifespan extension. This result clearly shows that these animals have enough stored resources for reproduction under DR and that suppression of reproduction is an adaptive decision to avoid producing offspring in a poor environment. Our findings contradict ‘resource allocation’, ‘nutrient recycling’, and ‘clean cupboards’ models of DR, as we further show that DR increases two measures post-DR fitness. These results fit the hypothesis that DR response evolves because of relaxed selection on somatic maintenance and survival in non-DR environments (Fig. 1). Because this hypothesis lacked formal theoretical foundation, we developed a new mathematical model for the evolution of the DR response via mutation accumulation (DRM)^20^, which demonstrates that the DR response can evolve under a broad range of conditions.

**Figure 1:**
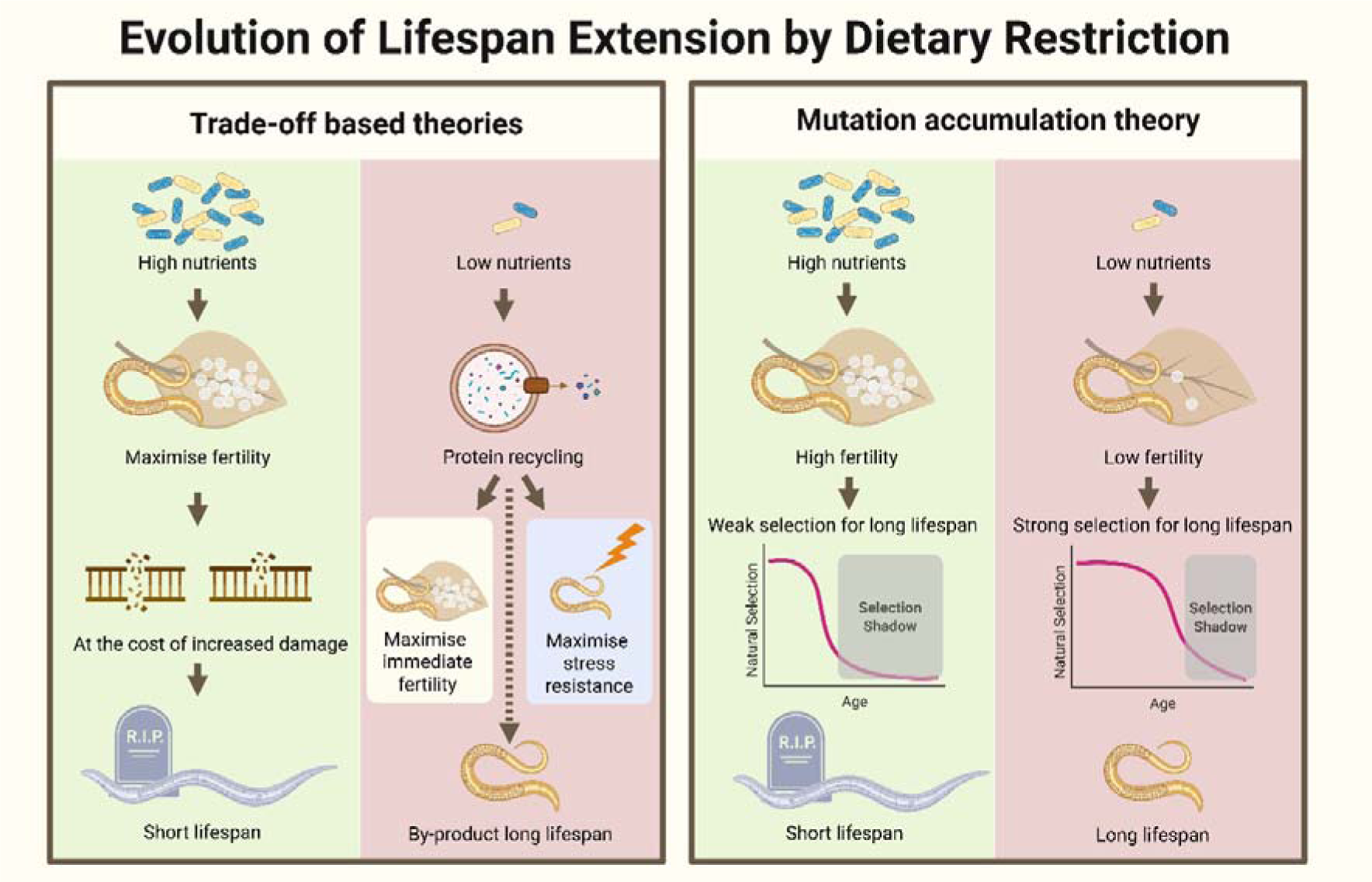
Theories for the evolution of lifespan extension by dietary restriction. Classical resource allocation trade-off theory maintains that increased fertility comes at the cost increased damage because fewer resources are allocated to repair when resources are plentiful. More recently, ‘recycling’ theories suggest that in low-nutrient environments, individuals recycle extraneous proteins to either maximise immediate fertility (‘nutrient recycling’) or immediate survival (‘clean cupboards’), leading to a longer lifespan as a by-product of toxic waste removal. Mutation accumulation theory maintains that reduced fertility under low nutrient availability leads to strong selection for longer lifespan in such environment, even in the absence of trade-offs, resulting in the evolution of dietary restriction response. This figure was created with BioRender.com.

## Results

### Empirical results

#### DR with or without food odour extends lifespan and improves stress resistance

Following treatments both with DR and DR+O, lifespan is significantly extended compared to *ad libitum* controls (Cox mixed effects model (Coxme): DR: Hazard ratio = 0.350, z = −5.94, *p* < 0.0001; DR+O: Hazard ratio = 0.350, z = −5.94, *p* < 0.0001, Fig. 2, Table S1), and adding odour has no significant effect on DR-based lifespan extension (Coxme Tukey post-hoc test: DR * DR+O: *p* = 0.999, Fig. 2, Table S1).

**Figure 2:**
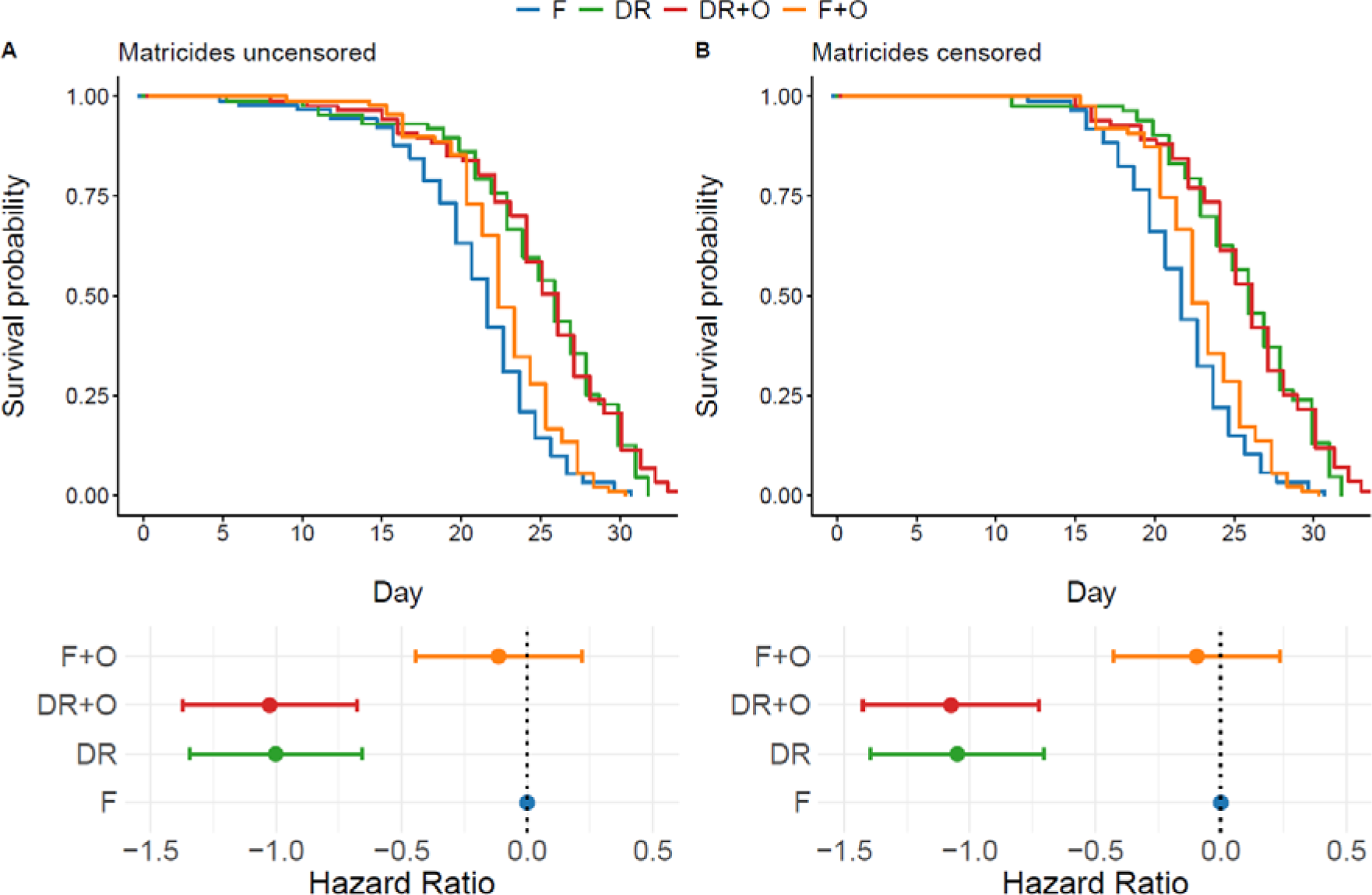
Dietary restriction extends lifespan with or without the presence of food odour. Survival probability and corresponding hazard ratios without matricides censored (A) and with matricides censored (B) following treatments of *ad libitum* food (F), *ad libitum* food plus food odour (F+O), dietary restriction (DR), and dietary restriction plus food odour (DR+O). Grey dotted lines represent baseline hazard ratio for F controls. Hazard ratios represented by means and 95% confidence intervals.

#### Food odour partially restores DR-driven fecundity decrease without cost to offspring

Dietary treatments significantly altered the reproductive schedule when compared to controls (Zero-inflated negative binomial (ZI NB) GLMM: Treatment*Day^2^: Χ^2^= 30.688, df = 3, *p* <0.0001; Treatment*Day: Χ^2^= 112.655, df = 3, p<0.0001, Table S2). DR+O had an intermediate effect on reproduction (DR+O*Day^2^: *p* = 0.0003; DR+O*Day: *p* < 0.0001, Table S2), slowing it down to a lesser extent than DR alone (DR* Day^2^: p <0.0001, DR*Day: p <0.0001, Fig. 3A). DR and DR+O treatments also significantly reduced lifetime reproductive success (LRS) (LM: R^2^ = 0.111, F_3,122_ = 6.219, p = 0.0006; DR: *p* = 0.003; DR+O: *p* = 0.017, Fig. 3B, Table S3). There was some evidence that the F+O treatment slightly altered the age-specific reproduction schedule (F+O*Day^2^: *p* = 0.003; F+O*Day: *p* = 0.028, Table S2), but this does not affect our conclusions.

**Figure 3:**
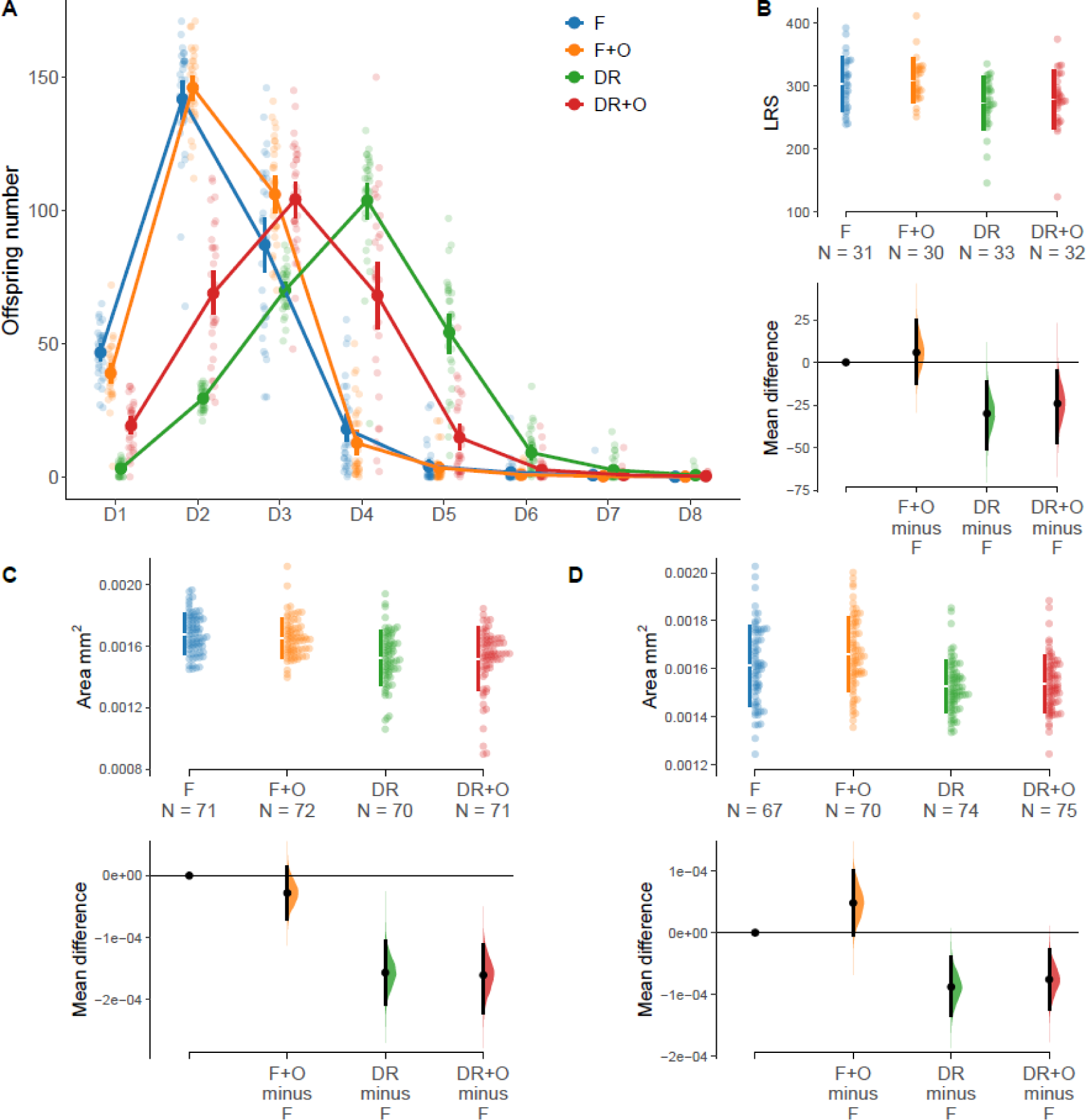
Dietary restriction slows reproduction, but to a lesser extent in the presence of food odour, without impacting egg size. Age-specific reproduction (A) and lifetime reproduction success (LRS; B) following treatments of *ad libitum* food (F), *ad libitum* food plus food odour (F+O), dietary restriction (DR), and dietary restriction plus food odour (DR+O). (A) points represent mean daily offspring production with 95% confidence intervals. (B) coloured points represent raw data with coloured lines to represent the standard deviation and the mean as the gap between these lines. Below this are bootstrapped mean differences and confidence intervals as compared to control (F) reference. Egg size (mm^2^) measured on day 2 (C) and day 4 (D) of adulthood. Coloured points represent raw data with coloured lines to represent the standard deviation and the mean as the gap between these lines. Below this are bootstrapped mean differences and confidence intervals as compared to control (F) reference.

DR and DR+O treatments also resulted in the production of smaller eggs than controls on Day 2 (LMM - *Χ*^2^= 27.262, df = 3, *p* < 0.0001; DR: *p* <0.001, DR+O: *p* <0.001, Fig. 3C, Table S4) and Day 4 (LMM - *Χ*^2^= 22.307, df = 3, *p* < 0.0001, DR: est. = *p* = 0.013, DR+O: *p* = 0.031, Fig. 3D, Table S4).

#### Mating restores lifetime reproductive success under DR with food odour exposure

Mated reproduction did not change the trends observed in age-specific reproduction observed in unmated hermaphrodites following dietary treatment (ZI NB GLMM: Treatment*Day: *Χ*^2^= 68.701, df = 3, *p* < 0.0001; Treatment*Day^2^: *Χ*^2^= 45.304, df = 3, *p* < 0.0001; Fig. 4A, Table S5). However, mating did allow LRS to return to control levels in DR+O treated individuals (LM: R^2^ = 0.176, F_3,67_ = 5.997, *p* = 0.001; DR+O: *p* = 0.565), but not for those treated with DR only (*p* = 0.002, Table S6, Fig. 4B).

**Figure 4:**
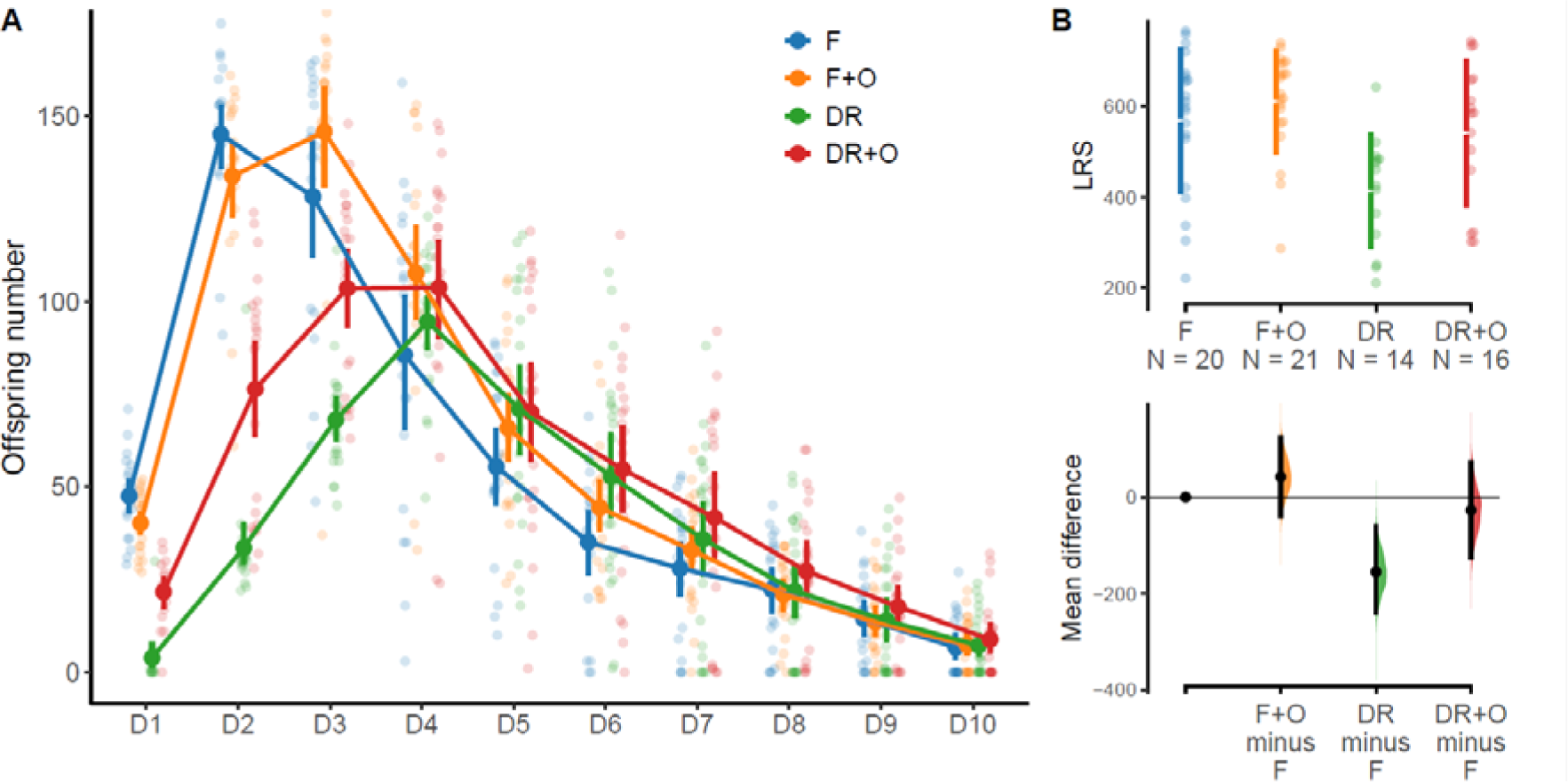
Mating restores lifetime reproductive success and partially restores individual fitness under dietary restriction with the presence of food odour. Mated age-specific reproduction (A) and lifetime reproduction success (LRS; B) following treatments of *ad libitum* food (F), *ad libitum* food plus food odour (F+O), dietary restriction (DR), and dietary restriction plus food odour (DR+O). (A) points represent mean daily offspring production with 95% confidence intervals. (B) coloured points represent raw data with coloured lines to represent the standard deviation and the mean as the gap between these lines. Below this are bootstrapped mean differences and confidence intervals as compared to control (F) reference.

#### Food odour under DR increases growth rates

DR and DR+O treatments result in smaller body size compared to controls on Days 2 (LM - R^2^ = 0.929, F_3, 92_ = 415, *p <* 0.0001; DR: est. *p* < 0.0001; DR+O: *p* <0.0001) and 4 (LM: R^2^ = 0.744, F_3, 92_ = 92.77, *p <* 0.0001; DR: *p* < 0.0001; DR+O: *p* < 0.0001, Fig. 5, Table S7). However, DR and DR+O treated individuals grew more than controls between Days 2 and 4 (LM: R^2^ = 0.503, F_3, 92_ = 33, *p <* 0.0001; DR: *p* < 0.0001; DR+O: *p* < 0.0001, Fig. 5, Table S7), and DR+O treatment induced faster growth than DR alone (LM post-hoc Tukey – DR*DR+O: p = 0.031, Table S8).

**Figure 5:**
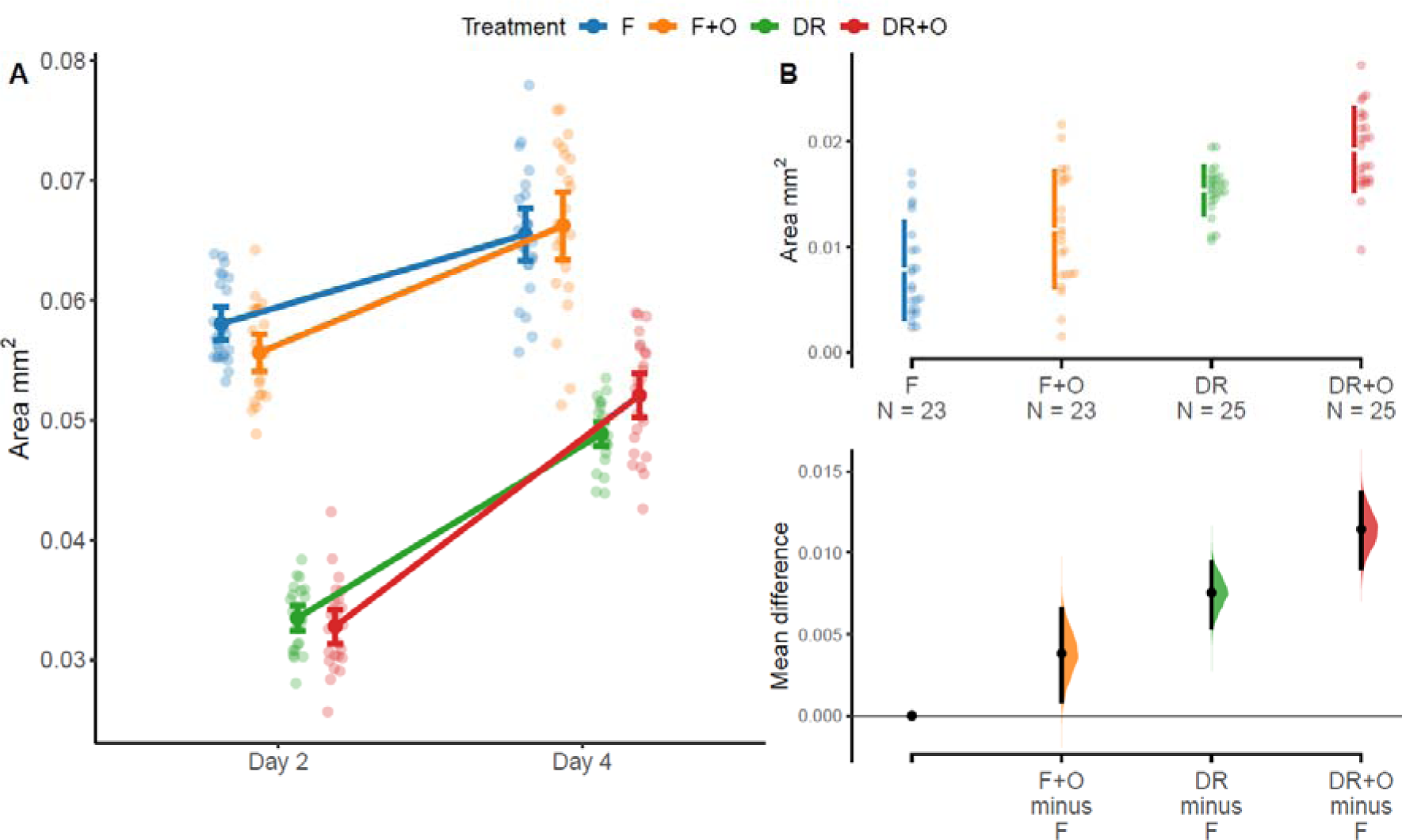
Presence of food odour under dietary restriction increases growth rates. Body size measured as surface area (mm^2^) on days 2 and 4 of adulthood (A) and growth rates (change in body surface area between days 2 and 4 of adulthood; B). (A) Dark points represent mean body size with 95% confidence intervals and transparent points represent raw data. (B) Coloured points represent raw data with coloured lines representing the standard deviation and the mean in the gap between these lines. Below this are bootstrapped mean differences and confidence intervals as compared to control (F) reference.

#### DR improves stress resistance and does not suppress population growth in semi-natural conditions

DR and DR+O treated individuals survived longer than controls (F) following heat shock (Coxme – DR: Hazard ratio: 0.042, se = 0.211, p < 0.0001; DR+O: Hazard ratio: 0.032, se = 0.214, p < 0.0001, Fig. 6 Table S9), and food odour under the *ad libitum* treatment had no effect on survival (Coxme – F+O: Hazard ratio: 1.097, se = 0.144, p = 0.522, Fig. 6, Table S9). In fact, all F and F+O individuals died before the end of the heat shock treatment; DR and DR+O nematodes survived 8- and 12-days post-heat shock, respectively.

**Figure 6:**
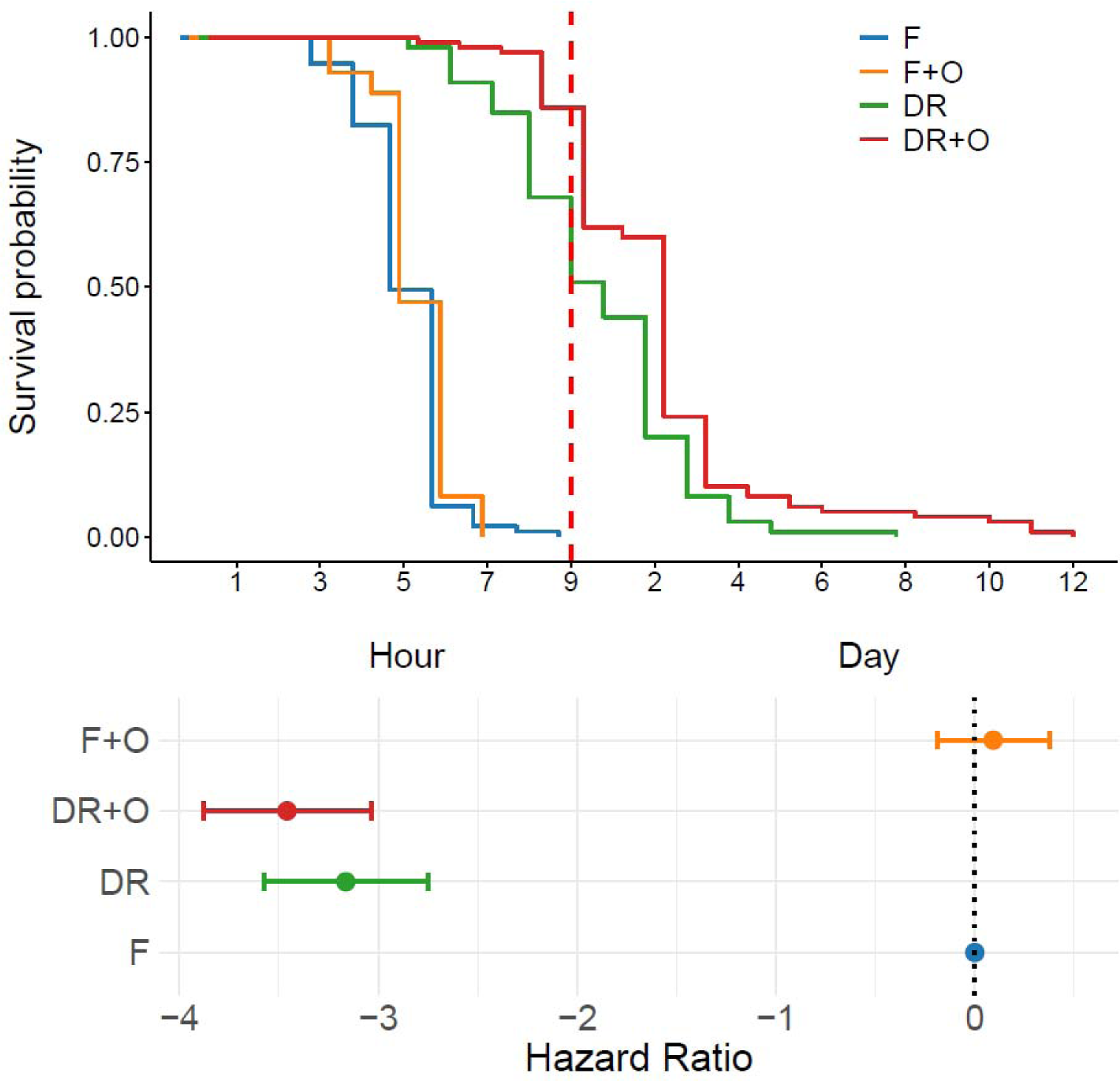
Dietary restriction with or without the presence of food odour improves resistance to heat stress. Survival probability and corresponding hazard ratios following treatments of *ad libitum* food (F), *ad libitum* food plus food odour (F+O), dietary restriction (DR), and dietary restriction plus food odour (DR+O). The r red dotted line represents the final hour of heat shock and the last hourly check for survival. Data to the right of this line were obtained through daily mortality checks following the heat shock. The grey dotted line represents baseline hazard ratio for F controls. Hazard ratios represented by means and 95% confidence intervals.

Our results do not indicate any cost to population growth following DR compared to fully fed controls. They point to a strong influence of environment on treatments, however, as is indicated by significant interactions with Block for all treatments across days and the quadratic variable for day (ZI NB GLMM: Treatment*Day: *Χ*^2^= 48.332, df = 6, *p* < 0.0001, Treatment*Day^2^: *Χ*^2^= 40.668, df = 6, *p* < 0.0001 Tables S10, S11, Fig. 7).

**Figure 7:**
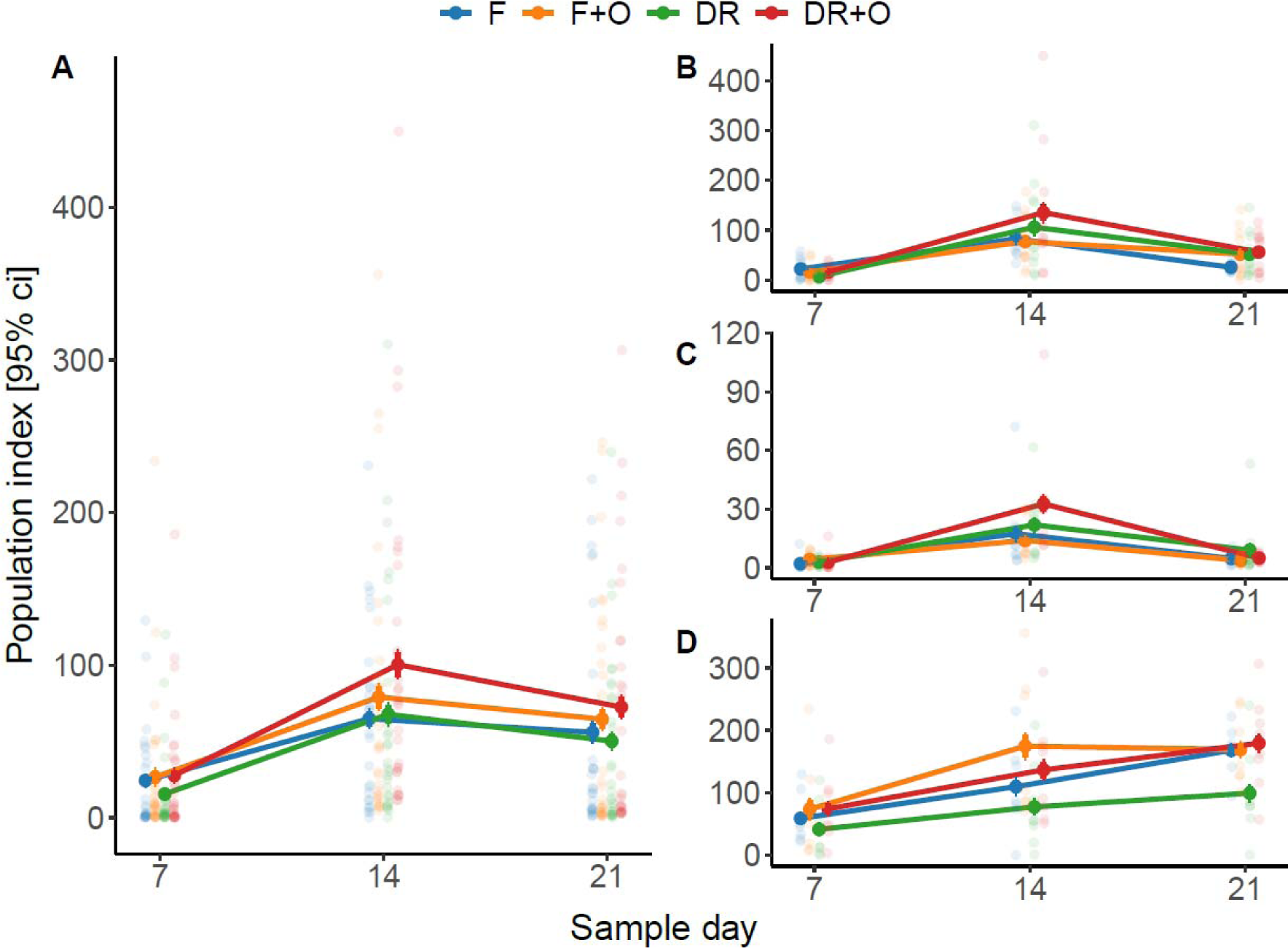
Early-life dietary restriction (DR) does not suppress post-DR population growth in semi-natural conditions. Population index, measured as the number of individuals counted in a compost sample from an outdoor microcosm population founded by 50 individuals following treatments of *ad libitum* food (F), *ad libitum* food plus food odour (F+O), dietary restriction (DR), and dietary restriction plus food odour (DR+O). (A) shows data across all blocks, and B, C and D show data from individuals blocks 1, 2, and 3, respectively. Transparent points represent raw data (mean number of individuals across six technical replicates per population), dark points represent means per treatment on a given day, and error bars show 95% confidence intervals.

### Evolutionary model

We imagine an age-structured population with overlapping generations that is exposed to a sequence of two different environments, E_1_, E_2_, which correspond to *ad libitum* and DR conditions, respectively. We assume that a move from E_l_ to E_2_ decreases age-specific fertility by proportion 1-k regardless of age. A shift from one environment to another will causes changes in the age structure that will take some time to equilibrate (stability). However, the population is free to grow or shrink while stable. The duration of each environmental exposure is assumed to be much longer than it takes to reach stability.

Fertility comes at a cost to survival, and this cost can vary with the environment. In *E*_l_, each reproductive event increases the immediate mortality rate of the asexual parent linearly in proportion to *c*-*I*; in *E*_2_ this is *c*+*I*. The added mortality realised in each environment follows from this per-unit cost and the rate of age-and-environment-specific fertility. The difference between realized costs of fertility is therefore *δ* = l(*k* +1) – *c*(1-*k*).

Finally, the population has genetic variation for age-specific mortality at all ages, and the genes that contribute to that variation are assumed to function in a strictly age-and-environment specific manner (i.e., there is no genetic correlation across mortality at different ages or across different environments). Genes that are specific to one environment are exposed to the effects of natural selection only when the population experiences that environment. At other times, these genes are neutral with respect to fitness.

The model that follows describes how fertility suppression experienced under DR (*E*_2_) shifts selection to enhance selection for increased adult lifespan slower senescence. Selection for age-specific mortality is derived using results from Hamilton (1966). Modelling details are presented in Supplement 1. A between-environment comparison of selection gradients finds conditions that favour the evolution of longer life in the fertility-suppressed environment (*E*_2_).

#### Case 1: No environmental differences in fertility costs, δ = 0

In this case, there are no fertility costs in either environment (*c*, *I* = 0) or, more generally, the realized fertility costs are the same in both environments, *k*(*c* + *I*) = *c*-*I*. Longevity selection is always enhanced in *E*_2_.

#### Case 2: There is a greater realized mortality cost to reproduction in *E*_2_, δ > 0

Specifically, δ > 0 if and only if *I* > *c*(1-*k*)⁄(1+*k*). These conditions always favor increased selection in *E*_2_.

The model is too complex to provide a more insightful general description of threshold conditions that define all cases. However, we can contrive of a simple case that demonstrates that conditions for enhanced lifespan selection can be satisfied when there are *less* realized mortality costs to reproduction in *E*_2_ (δ < 0).

#### Case 3: There is a less realized mortality cost to reproduction in *E*_2_, δ < 0

Enhanced selection for adult lifespan and slower senescence in *E*_2_ is achieved when the differences in realized fertility costs are on the interval 0 > δ > -(1-*k*). In terms of per-offspring costs, δ = *I*(*k*+ 1) - *c*(1-*k*). For example, this condition is met when fertility is suppressed by 50% in *E*_2_, and each offspring increases age-specific mortality by 1.5 and 2.5 in *E*_l_ and *E*_2_, respectively.

#### Evolution of a plastic lifespan response

We have made two sets of model assumptions. The first involves genetic architecture (age-and-environment-specific gene action), and the second involves environmental and physiological conditions (fertility suppression and costs to reproduction). When combined, our model predicts the evolution of among-environment variation in adult lifespan and actuarial senescence between environments that represent *ad libitum* and DR conditions; this is an evolved plastic response. The predictions apply to a broad range of parameter values. Perhaps the most compelling is the prediction that fertility suppression by itself (Case 1) is sufficient to drive the evolution of divergent lifespan. When mortality costs of reproduction are added to the model, the predictions become more complex. Costs that cause a net increase in mortality in the DR environment (Case 2) always lead to the evolution of lifespan extension in this environment, but when the net costs of reproduction in the DR are less than in the *ad libitum* case, then the predictions because more nuanced. Nevertheless, we have demonstrated that cases exist where the DR lifespan response can evolve when costs under DR are low (Case 3).

#### Population stability

The model of selection provided by Hamilton^15^ assumes stable age-distributions. As noted above, this requires that the durations of each environment are long. This feature has positive and negative attributes. On the one hand, this reveals that the DR lifespan response can evolve when individuals never experience more than one environment. In fact, the greater the proportion of individuals that *don’t* experience both environments, the better this assumption is met. This prediction is novel, as other models imply that the selection that leads to the evolution of the DR response are driven by individuals waiting out bad times in hopes of living long enough to experience good times. On the other hand, our model cannot cope directly with environmental fluctuations that are rapid (when compared to the lives of the individuals). However, we can describe how the models break-down and make a reasonable inference about evolutionary outcomes in these cases.

The evolution of the plastic DR response described in our models is predicated on divergent selection regimes that are driven directly by differences in the age structure of new parents (the integrand in S1 defines the probability distribution function of these parents). These structural differences emerge over time as the age structure equilibrates after an environmental shift; it is only at the demographic equilibria where Hamilton’s expressions are strictly correct. However, it is reasonable to expect that as the age-structure shifts from the equilibrium state at one environment to that of another, selection shifts from the one deriving from the first equilibrium to the one deriving from the second. While selection is in flux, it is likely to reflect a character that is intermediate to selection regimes defined at both equilibrium states. It is also reasonable to expect that as the duration of each environment shortens, this intermediate selection regime operates at a greater proportion of time. If selection becomes more intermediate over time, then it must follow that the magnitude of the plastic lifespan response will evolve to be less and less. Differently put, the potential for this mechanism to drive the evolution of the DR response follows inversely to the frequency of environmental shifts.

The model that we described proposes an evolutionary pathway to extended lifespan under DR conditions that operates in a fundamentally different way and under different conditions than those previously proposed. In addition, it is the first formal evolutionary model of the phenomenon that leverages Hamilton’s description of selection gradients, a cornerstone of the modern evolutionary theory of ageing^21–23^.

## Discussion

The evolution of lifespan extension under DR has been thought either to be driven by reallocation of resources from reproduction to somatic maintenance, or as a by-product of nutrient recycling. Our empirical results challenge both suppositions. We show that DR animals can maintain stored resources that would allow them to increase their reproductive rate, up to a point, without any effect on lifespan. Importantly, DR supresses reproductive rate, also in the presence of food odour, even when total reproduction is unaffected. We present a new evolutionary model that demonstrates that the DR-driven lifespan extension can evolve via a route that does not require resource re-allocation or other trade-offs (Fig. 1). Instead, the model assumes that DR suppresses reproduction, and this leads to enhanced selection for long life.

If genes are expressed differently in DR and non-DR environments, then this change in natural selection can be sufficient for the evolution of increased somatic maintenance in DR environments. Such differential gene expression is commonly reported in genomic studies of DR^24–26^. This parsimonious model is rooted in standard evolutionary theory, and reconciles empirical findings presented here with the previous studies. This model suggests that lifespan extension under DR evolves under a broader range of conditions than has been previously considered, namely a relaxation of the previous assumption that individual organisms experience both environments during their lifetime. Furthermore, because trade-offs are unnecessary for this model to work, the DR response can evolve via germline mutation accumulation consistent with the Mutation Accumulation model of ageing^20^. Importantly, this model is in line with experimental evolution studies suggesting that evolution of fertility and lifespan under DR can be uncoupled^27,28^.

While this DR-driven longevity via mutation accumulation model (DRM) does not require any trade-offs for increased somatic maintenance under DR to evolve, it does not preclude such trade-offs. In fact, the existence of genetic trade-offs would make it easier to evolve longer lifespan under DR. Therefore, different forms of genetic trade-offs represent special cases of a general model, while the only overarching assumption is that DR reduces fertility.

### To reproduce, or not to reproduce

Disentangling the physiological versus selection-based effects of resource limitation on reproduction was possible in our experiment by engaging both gustatory and olfactory responses in *C. elegans.* By providing the odour of the food, but without actual food available to eat, only the nematodes’ olfactory responses were engaged, which was enough to induce reproduction, albeit at a lower rate than fully fed individuals. This indicates that although DR individuals carry enough resources to produce some offspring (whether this is through recycling extraneous cellular and protein materials or reserves) they reproduce less when they sense that their environment is poor. Even when reproduction is induced in the presence of food odour without access to food, there does not appear to be any trade-off between reproduction and somatic maintenance. While another study found that individuals that reproduced in the absence of food did not experience the lifespan extension indicative of a trade-off^19^, their treatment was 48 hours of starvation compared to our 24-hour treatment, highlighting that the line between DR and malnutrition can be difficult to discern.

Our results also contradict the notion that the DR response is solely a laboratory artifact. It is known that the *E. coli* fed to laboratory *C. elegans* is damaging, especially at older ages, and shortens lifespan^29^, but our transient treatment ensures that the effects on nematodes subjected to DR cannot be due to avoidance of such effects. Furthermore, our study design proves that the delay in reproduction is decision-based, rather than a consequence of limited resource availability. Resource availability plays an integral role in determining the life cycle of *C. elegans* during development^30^, and it is likely that selection has also acted on response to food availability during reproductive stages. Furthermore, some suggest that DR response encompasses much more than response to nutrient availability, which is likely only a part of a response to conditions expected to coincide with resource scarcity in nature, such as heatwaves or drought^31^. This is consistent with our results and those of others who show that nutrient restriction improves resistance to stress^13^. Improved stress resistance may explain the results we see in our semi-natural populations following DR and DR+O treatment. While DR and DR+O worms reproduce more slowly, improved resistance to stress may compensate for this compared to our control treatments, allowing post-DR populations to grow at similar rates and DR+O populations to grow faster.

### Broad implications of the model

While we focused here on the reasons behind DR-driven lifespan extension, it is important to note that this model is broadly applicable to any kind of environmental stress that reduces fertility. The evolution of hormesis, defined here as non-lethal stress that increases somatic maintenance and lifespan, is a long-standing problem in biology. The model presented here suggests that any kind of non-lethal stress that reduces reproduction will result in selection favouring increased survival, and therefore, increased somatic maintenance. This process suggests that hormesis is expected to evolve under a very broad range of conditions, because reduced fertility often accompanies non-lethal stress. For example, elegant work on germline-soma interactions in *C. elegans* showed that when the germline is damaged by the environmental stress and needs time to recover, it upregulates systemic stress response in the somatic cells resulting in increased survival and stress resistance^32^. This example fits perfectly with the model presented here, because extrinsic damage to germline reduces fertility and changes selection gradients for survival. The model suggests that selection on upregulation of innate immune response and increased somatic endurance when germline integrity is temporarily compromised is stronger than when the germline is intact, and fertility is high.

## Conclusions

Our results and model demonstrate that previous trade-off-based models are neither sufficient nor necessary to explain lifespan extension under DR in the general case. We show that germline mutation accumulation could lead to the evolution of shorter lifespans in environments where nutrients are plentiful. With the plethora of conflicting results across species, genotypic backgrounds, and laboratory techniques throughout the field, it is no surprise that both the evolution and mechanisms of DR remain hotly debated. We suggest that our model provides a general and parsimonious explanation for the evolution of increased somatic maintenance and lifespan extension under DR. Importantly, this model can be used to predict the evolution of longer lifespan under any environmental stress that supresses fertility and, therefore, changes selection gradients for adult survival in any organism.

## Methods

### Experimental model

N2 wild-type *Caenorhabditis elegans* nematodes were obtained from the *Caenorhabditis* Genetics Centre (funded by NIH Office of Research Infrastructure Programs - P40 OD010440). Populations maintained in −70°C were thawed, then bleached using a solution of NaOCl and NaOH to rid the population of fungal and bacterial contamination. To prevent effects of freezing from impacting experimental work, populations were maintained for at least two generations on 90 mm Nematode Growth Medium (NGM) agar plates made with 10 mg ml^-1^ of Nystatin and 100 mg ml^-1^ of ampicillin at 20°C. Plates were seeded with *Escherichia coli* OP50-1 (pUC4 K), obtained from J. Ewbank at the Centre d’Immunologie de Marseille-Luminy, France to feed the populations. Short-duration timed egg lays were used to produce age-synchronised populations.

### Experimental design

The experiments described here consisted of four treatments: 1) *ad libitum* food (F); 2) *ad libitum* food plus food odour (F+O); 3) dietary restriction via 24-hours without access to food (DR); and 4) dietary restriction plus food odour (DR+O). The control treatment (F) consisted of NGM agar plates seeded with a lawn of *E. coli.* DR plates were made without peptone, which is included in NGM agar plates to allow bacterial growth and were left unseeded. DR+O and F+O plates contained two agar layers, both with a bottom layer of NGM agar seeded with *E. coli* that was allowed to grow overnight. F+O plates then had a second layer identical to the bottom layer. DR+O plates had a second layer made of NGM agar without peptone and were left unseeded. Bacterial incubation times were standardised across treatments.

For all experiments described here, age-synchronised populations of nematodes were produced via egg lay of Day 2 adults. Upon reaching late L4 stage, nematodes were placed on their treatment of F, F+O, DR, or DR+O for a 24-hour period. Excess individuals were placed onto DR and DR+O treatments due to high risk of unfed individuals crawling up the side of the plastic plates and dying. When more than the desired number of individuals remained after treatment, the necessary number of individuals were randomly selected for use in the assay.

### Lifespan and reproduction assays

First, the effects of DR and DR+O on reproduction and longevity were tested in the laboratory, along with F controls and F+O positive controls. Individuals were kept on their own 35 mm NGM plates for reproduction assays, and lifespan assays consisted of 35 mm NGM plates containing 10 individuals. To determine daily survival, observations of touch-response were used.

To test reproductive effects of DR and DR+O, individuals from all treatments were transferred to new plates every 24 hours. Eggs obtained each day were reared for two days, then killed by placing into a 42°C incubator for 3.5 hours in order to count them. Assays were fully blinded following the initial treatment day. Reproduction assays continued until approximately 75 percent of individuals stopped reproducing for at least one day.

### Mated reproduction assay

To identify what role sperm limitation in hermaphrodites may play in reduced reproduction following DR, non-experimental males were introduced to hermaphrodites for 24 hours. Reproduction after mating was measured using the methods described above. Mating took place immediately following treatment, on Day 2 of adulthood. Mating had to be performed after treatment to avoid treatment effects on males impacting the experiment. Each reproductive assay plate held 1 hermaphrodite and 2 males for 24-hours. The males were removed upon transfer to Day 3 reproductive assay plates, and the assay resumed as described above.

### Body and egg size assays

Following treatment, on Days 2 and 4 of adulthood, individuals were placed on fresh plates and immediately photographed using a Leica MC170 HD camera attached to a Leice M165 C light microscope. The individuals were then allowed to lay eggs for 2-4 hours. After, up to 3 eggs were randomly chosen to take images of. Days 2 and 4 were chosen according to the reproductive peaks of fully fed worms (Day 2) and starved worms (Day 4). Surface area measurements were taken in Imagej^33^ by turning colour images into binary images, filling in transparent areas in individuals not captured by the automated binary tool in ImageJ, and measuring the black areas. The same method was used to measure eggs.

### Heat shock assay

The heat shock assay was performed similarly to a lifespan experiment. Following exposure to their respective treatments, nematodes were placed onto fresh seeded plates in groups of 10. The plates were then placed in a 37 °C incubator and checked hourly for response to touch. As heat shock can cause immobility in nematodes, those assumed dead were kept on the plate in a designated location to confirm mortality at subsequent checks. Heat shock lasted 9 hours, after which surviving nematodes were kept at 20°C and monitored for survival over the following days until all individuals died.

### Outdoor assay

Outdoor populations were housed in organic, peat-free compost in 250-ml conical tubes. The tubes had holes in the bottom to allow for drainage in the case of excess rain. The tubes were suspended above a bath of heavily salted water to kill any nematodes that escape through those holes to prevent environmental contamination. Compost was autoclaved no more than 7 days before the start of the experiment, with OP50 *E. coli* (susceptible to antibiotics) being reintroduced to the compost as a food source for the populations. The OP50 was maintained in LB broth at 4°C after growing at 37°C for 18 hours. The compost was completely saturated in OP50-inoculated LB broth (60 ml) and allowed to grow at room temperature for 24 hours before nematodes were introduced.

The F, F+O, DR, and DR+O treatments were given on 90 mm agar plates for this experiment, due to the high number of nematodes required. Following treatment, 50 individuals were collected from a given treatment and placed on a clean 35-mm agar plate, then rinsed into a tube of compost using OP50 bacterial broth. Outdoor populations were fed weekly with 25 ml of OP50.

Outdoor populations were sampled weekly over a 3-week period. Each week, a 5-ml sample of compost was retrieved from each tube, after mixing the top half of the compost in each conical tube. Samples were cleaned with osmosis purified (RO) water to remove small particles by dissolving the compost in RO water at a total volume of 10 ml and centrifuging at 1800 g for 2 minutes, then pouring out the supernatant. Following this cleaning, the compost was then dissolved in a MgSO_4_ solution with a specific density of 1.15 (at which nematodes will be suspended) at a total volume of 10 ml and again centrifuged at 1800 g for 2 minutes. The supernatant was then carefully mixed with a pipette, then 0.1-ml samples of the supernatant were pipetted onto 35-mm agar plates and allowed to dry for future counting. For each population, 6 technical replicate samples were taken.

### Statistical analysis

All analyses were performed in R version 4.4.1^34^. Model diagnostics were performed with the DHARMa package^35^. Where overdispersion was present in generalised linear mixed models, observation level random effects were included to control for it. Model selection was decided by choosing models with the lowest Akaike information criterion (AIC) based on error structures, zero-inflation parameters, and dispersion parameters. Bootstrap-coupled estimation plots were produced with the dabestr package^36^, and all other plots were created using ggplot2^37^.

Survival assays were analysed by using cox proportional hazards models with mixed effects from the coxme^38^ and survival^39^ packages. These models fit Treatment as a fixed effect, and Plate ID as a random intercept. This model was performed with and without matricides. Heat shock survival data was also analysed using these models with the same parameters.

Age-specific reproduction was analysed using zero-inflated negative binomial generalised linear mixed models (GLMMs) with the glmmTMB package^40^ with observation level random effects, including interacting fixed factors of Treatment, Day^2^, and Day and individual ID as a random intercept to account for repeated measures of individuals reproductive output over time. The full model for unmated age-specific reproduction is as follows: No_offspring ∼ Treatment * poly(Day, 2) + (1|ID), zi = ∼ Day, disp = ∼Day. For mated reproduction the final model is as follows: No_offspring ∼ Treatment * poly(Day, 2) + (1|ID), zi = ∼ poly(Day,2), disp = ∼Treatment*Day^2^.

The effects of dietary treatment on LRS and individual fitness and body size were analysed using linear models with Gaussian error structures were used, and to identify differences between treatments, the *emmeans* package^41^ was used to perform pairwise comparisons. The effect of dietary treatment on egg size was analysed using linear mixed effects models with Gaussian error structures with Treatment as a fixed factor and ID as a random intercept to account for repeated measures of technical replicates per individual.

Population growth data was analysed using zero-inflated negative binomial GLMMs with the glmmTMB package. Treatment, Day^2^, Day and Block were fit as fixed factors, with all interactions in initial model. The final model is as follows: Population index ∼ Treatment* poly(Day, 2) *Block. Technical replicate nested within Population was included as a random intercept to account for repeated measures within populations across replicates and time. To account for problems with overdispersion and zero inflation, formulas were fit for each. The final model includes a ziformula = ∼ Day + Block.

## Supporting information

Supplementary Data Analysis and Model Materials

## Data availability

All code and data used for this study are available via GitHub at https://github.com/irishsar92/DR_MAT.

## Notes

https://github.com/irishsar92/DR_MAT

### Competing Interest Statement

The authors have declared no competing interest.

### Summary of Updates

Figure 1 legend revised; Figure 2 revised

