## Supplementary Data Analysis and Model Materials for "Mutation accumulation underpins evolution of lifespan extension by dietary restriction"

**Supplementary Data Analysis Materials:**

**Table S1:** Cox mixed effects model summary for nematode survival data following treatments of *ad libitum* food (F), *ad libitum* food plus odour (F+O), dietary restriction (24-hour starvation; DR), or dietary restriction plus odour (DR+O). Model was performed on data including and excluding matricides as deaths, and results from data censoring matricides is used for interpretation and post-hoc analysis.

| **Matricides included** | | | | | | | | | |
| --- | --- | --- | --- | --- | --- | --- | --- | --- | --- |
|  | *Coefficient* | | *Exp(Coefficient)* | | *Std. Error* | | *z* | | *p* |
| DR | -1.0014 | | 0.3674 | | 0.1755 | | -5.71 | | <0.0001 |
| DR+O | -1.0254 | | 0.3587 | | 0.1775 | | -5.78 | | <0.0001 |
| F+O | -0.1141 | | 0.8922 | | 0.1689 | | -0.68 | | 0.499 |
| Random effects: Plate.ID | | | Variance: 0.0282 | | | | Std. Deviation: 0.1682 | | |
| **Matricides censored** | | | | | | | | | |
| DR | -1.0494 | | 0.3502 | | 0.1766 | | -5.94 | | <0.0001 |
| DR+O | -1.0745 | | 0.3415 | | 0.1788 | | -6.01 | | <0.0001 |
| F+O | -0.9579 | | 0.9087 | | 0.1690 | | -0.57 | | 0.571 |
| Random effects: Plate.ID | | | Variance: 0.0244 | | | | Std. Deviation: 0.1561 | | |
| **Tukey post-hoc comparisons (Matricides censored)** | | | | | | | | | |
|  | | *Estimate* | | *Std. Error* | | *z* | | *p* | |
| F – DR | | 1.094 | | 0.177 | | 5.943 | | <0.0001 | |
| F – DR+O | | 1.075 | | 0.179 | | 6.009 | | <0.0001 | |
| F – F+O | | 0.096 | | 0.169 | | 0.567 | | 0.942 | |
| DR – DR+O | | 0.025 | | 0.173 | | 0.145 | | 0.999 | |
| DR – F+O | | -0.954 | | 0.176 | | -5.403 | | <0.0001 | |
| DR+O – F+O | | -0.979 | | 0.179 | | -5.476 | | <0.0001 | |

**Table S2:** Zero-inflated negative binomial generalised linear mixed effects model results for analysis of age-specific reproduction over eight days in nematodes following treatments of *ad libitum* food (F), *ad libitum* food plus odour (F+O), dietary restriction (24-hour starvation; DR), or dietary restriction plus odour (DR+O).

| **Model: No_offspring ~ Treatment*Day + Treatment*Day^2 + (1\|ID), zi = ~ Treatment** | | | | | | | | |
| --- | --- | --- | --- | --- | --- | --- | --- | --- |
|  | *Estimate* | | *Std. Error* | | *z-value* | | | *p-value* |
| (Intercept) | 3.928 | | 0.267 | | 14.705 | | | <0.0001 |
| F+O | -0.616 | | 0.394 | | -1.563 | | | 0.118 |
| DR | -4.826 | | 0.355 | | -13.579 | | | <0.0001 |
| DR+O | -2.179 | | 0.355 | | -6.133 | | | <0.0001 |
| Day | 0.719 | | 0.162 | | 4.436 | | | <0.0001 |
| Day^2^ | -0.214 | | 0.021 | | -9.984 | | | <0.0001 |
| F+O: Day | 0.543 | | 0.246 | | 2.204 | | | 0.028 |
| DR: Day | 2.045 | | 0.203 | | 10.052 | | | <0.0001 |
| DR+O: Day | 1.094 | | 0.211 | | 5.181 | | | <0.0001 |
| F+O: Day^2^ | -0.101 | | 0.034 | | -2.954 | | | 0.003 |
| DR: Day^2^ | -0.143 | | 0.026 | | -5.536 | | | <0.0001 |
| DR+O: Day^2^ | -0.096 | | 0.028 | | -3.455 | | | 0.0006 |
| **Zero-inflation formula = ~ Treatment** | | | | | | | | |
| (Intercept) | -2.906 | | 0.504 | | -5.759 | | | <0.0001 |
| F+O | -0.752 | | 1.006 | | -0.748 | | | 0.455 |
| DR | -4.102 | | 14.182 | | -0.289 | | | 0.772 |
| DR+O | -1.299 | | 1.078 | | -1.205 | | | 0.228 |
| Random effects: ID | | | Variance: 0.130 | | | Std. Deviation: 0.360 | | |
| **Type III Anova** | | |  | | |  | | |
|  | | $\chi^{2}$ | | *df* | | | *p* | |
| (Intercept) | | 216.250 | | 1 | | | <0.0001 | |
| Treatment | | 220.297 | | 3 | | | <0.0001 | |
| Day | | 19.678 | | 1 | | | <0.0001 | |
| Day^2^ | | 99.683 | | 1 | | | <0.0001 | |
| Treatment*Day | | 112.655 | | 3 | | | <0.0001 | |
| Treatment* Day^2^ | | 30.688 | | 3 | | | <0.0001 | |

**Table S3:** Linear model output from analysis of lifetime reproductive success (LRS, total number of offspring produced) in nematodes following treatments of *ad libitum* food (F), *ad libitum* food plus odour (F+O), dietary restriction (24-hour starvation; DR), or dietary restriction plus odour (DR+O).

| **Model: LRS ~ Treatment:** R^2^ = 0.111, F_3,122_ = 6.219, p = 0.0006 | | | | |
| --- | --- | --- | --- | --- |
|  | *Estimate* | *Std. Error* | *t-value* | *p-value* |
| (Intercept) | 302.935 | 7.136 | 42.453 | <0.0001 |
| F+O | 5.765 | 10.175 | 0.567 | 0.572 |
| DR | -30.087 | 9.937 | -3.028 | 0.003 |
| DR+O | -24.217 | 10.012 | -2.419 | 0.017 |

**Table S4:** Linear mixed effects model output for analysis of egg size on Days 2 and 4 of adulthood in nematodes following Day 1 treatment of *ad libitum* food (F), *ad libitum* food plus odour (F+O), dietary restriction (24-hour starvation; DR), or dietary restriction plus odour (DR+O).

| **Model: Day 2 Egg Size ~ Treatment + (1\|ID):** *Χ^2^=* 27.262, df = 3, *p* < 0.0001 | | | | | |
| --- | --- | --- | --- | --- | --- |
|  | *Estimate* | *Std. Error* | *df* | *t-value* | *p-value* |
| (Intercept) | 1.677*10^-3^ | 2.813*10^-3^ | 91.77 | 59.616 | <0.0001 |
| F+O | -2.609*10^-5^ | 3.974*10^-3^ | 91.47 | -0.657 | 0.513 |
| DR | -1.569*10^-4^ | 3.981*10^-3^ | 92.07 | -3.941 | <0.001 |
| DR+O | -1.606*10^-4^ | 3.978*10^-3^ | 91.77 | -4.037 | <0.001 |
| Random effects: ID | | 1.545*10^-8^ | Std. Deviation | | 1.243*10^-4^ |
| Residual variance: | | 1.041*10^-8^ | Std. Deviation | | 1.020*10^-4^ |
| **Model: Day 4 Egg Size ~ Treatment + (1\|ID):** *Χ^2^=* 22.307, df = 3, *p* < 0.0001 | | | | | |
| (Intercept) | 1.608*10^-3^ | 2.312*10^-5^ | 99.88 | 69.540 | <0.0001 |
| F+O | 5.186*10^-5^ | 3.288*10^-5^ | 94.3 | 1.577 | 0.118 |
| DR | -8.073*10^-5^ | 3.187*10^-5^ | 105.50 | -2.533 | 0.013 |
| DR+O | -7.125*10^-5^ | 3.248*10^-5^ | 93.71 | -2.194 | 0.031 |
| Random effects: ID | | 9.999*10^-9^ | Std. Deviation | | 1.000*10^-4^ |
| Residual variance: | | 9.015*10^-9^ | Std. Deviation | | 9.495*10^-5^ |

**Table S5:** Zero-inflated negative binomial generalised linear mixed effects model results for analysis of age-specific reproduction over ten days in mated nematodes following treatments of *ad libitum* food (F), *ad libitum* food plus odour (F+O), dietary restriction (24-hour starvation; DR), or dietary restriction plus odour (DR+O).

| **Model: No_offspring ~ Treatment * poly(Day, 2) + (1\|ID), zi = ~ Day** | | | | | | | | | |
| --- | --- | --- | --- | --- | --- | --- | --- | --- | --- |
|  | | *Estimate* | | *Std. Error* | | *z-value* | | | *p-value* |
| (Intercept) | | 4.597 | | 0.172 | | 26.777 | | | <0.0001 |
| F+O | | -0.317 | | 0.236 | | -1.340 | | | 0.180 |
| DR | | -2.531 | | 0.269 | | -9.421 | | | <0.0001 |
| DR+O | | -1.127 | | 0.248 | | -4.550 | | | <0.0001 |
| Day | | 0.055 | | 0.073 | | 0.761 | | | 0.447 |
| Day^2^ | | -0.028 | | 0.007 | | -4.331 | | | 0.0001 |
| F+O: Day | | 0.190 | | 0.099 | | 1.929 | | | 0.054 |
| DR: Day | | 0.849 | | 0.108 | | 7.861 | | | <0.0001 |
| DR+O: Day | | 0.421 | | 0.102 | | 4.144 | | | 0.0003 |
| F+O: Day^2^ | | -0.018 | | 0.009 | | -2.065 | | | 0.039 |
| DR: Day^2^ | | -0.062 | | 0.009 | | -6.524 | | | <0.0001 |
| DR+O: Day^2^ | | -0.031 | | 0.009 | | -3.413 | | | 0.0006 |
| **Zero-inflation formula = ~Treatment*Day** | | | | | | | | | |
| (Intercept) | | -7.525 | | 1.408 | | -5.344 | | | <0.0001 |
| F+O | | -4.333 | | 3.330 | | -1.301 | | | 0.193 |
| DR | | 3.971 | | 1.539 | | 2.580 | | | 0.010 |
| DR+O | | 0.965 | | 1.899 | | 0.508 | | | 0.611 |
| Day^2^ | | 0.746 | | 0.163 | | 4.574 | | | <0.0001 |
| F+O: Day^2^ | | 0.382 | | 0.364 | | 1.049 | | | 0.294 |
| DR: Day^2^ | | -0.470 | | 0.183 | | -2.569 | | | 0.010 |
| DR+O: Day^2^ | | -0.161 | | 0.221 | | -0.730 | | | 0.465 |
| Random effects: ID | | | | Variance: 0.036 | | | Std. Deviation: 0.189 | | |
| Type III Anova | | | | | | | | | |
|  | | | $\chi^{2}$ | | *df* | | | *p* | |
| (Intercept) | | | 717.008 | | 1 | | | <0.0001 | |
| Treatment | | | 103.841 | | 3 | | | <0.0001 | |
| Day | | | 0.579 | | 1 | | | 0.447 | |
| Day^2^ | | | 18.755 | | 1 | | | 0.0001 | |
| Treatment*Day | | | 68.701 | | 3 | | | <0.0001 | |
| Treatment*Day^2^ | | | 45.304 | | 3 | | | <0.0001 | |

**Table S6:** Linear model output from analysis of lifetime reproductive success (LRS, total number of offspring produced) in mated nematodes following treatments of *ad libitum* food (F), *ad libitum* food plus odour (F+O), dietary restriction (24-hour starvation; DR), or dietary restriction plus odour (DR+O).

| **Model: Mated LRS ~ Treatment:** R^2^ = 0.176, F_3,67_ = 5.997, *p* = 0.001 | | | | |
| --- | --- | --- | --- | --- |
|  | *Estimate* | *Std. Error* | *t-value* | *p-value* |
| (Intercept) | 568.45 | 31.01 | 18.333 | <0.0001 |
| F+O | 42.79 | 43.33 | 0.988 | 0.327 |
| DR | -155.09 | 48.32 | -3.210 | 0.002 |
| DR+O | -26.89 | 46.51 | -0.578 | 0.565 |

**Table S7:** Linear model output for analysis of adult nematode body size on Days 2 and 4 of adulthood and growth between Days 2 and 4 following treatment on Day 1 of adulthood of *ad libitum* food (F), *ad libitum* food plus odour (F+O), dietary restriction (24-hour starvation; DR), or dietary restriction plus odour (DR+O).

| **Model: Day 2 Body Size ~ Treatment:** R^2^ = 0.929, F_3, 92_ = 415, *p <* 0.0001 | | | | |
| --- | --- | --- | --- | --- |
|  | *Estimate* | *Std. Error* | *t-value* | *p-value* |
| (Intercept) | 0.0581 | 0.0007 | 84.429 | <0.0001 |
| F+O | -0.0024 | 0.0010 | -2.505 | 0.014 |
| DR | -0.0246 | 0.0010 | -25.517 | <0.0001 |
| DR+O | -0.0253 | 0.0010 | -26.232 | <0.0001 |
| **Model: Day 4 Body Size ~ Treatment:** R^2^ = 0.744, F_3, 92_ = 92.77, *p <* 0.0001 | | | | |
| (Intercept) | 0.0655 | 0.0010 | 63.584 | <0.0001 |
| F+O | 0.0007 | 0.0015 | 0.502 | 0.617 |
| DR | -0.0166 | 0.0014 | -11.538 | <0.0001 |
| DR+O | -0.0134 | 0.0014 | -9.318 | <0.0001 |
| **Model: (D4 Size–D2 Size) ~ Treatment:** R^2^ = 0.503, F_3, 92_ = 33, *p <* 0.0001 | | | | |
| (Intercept) | 0.0074 | 0.0010 | 7.422 | <0.0001 |
| F+O | 0.0032 | 0.0014 | 2.232 | 0.028 |
| DR | 0.0079 | 0.0014 | 5.643 | <0.0001 |
| DR+O | 0.0118 | 0.0014 | 8.413 | <0.0001 |

**Table S8:** Post-hoc analysis using Tukey method of linear model analysis of growth between Days 2 and 4 of adulthood in nematodes following treatments of *ad libitum* food (F), *ad libitum* food plus odour (F+O), dietary restriction (24-hour starvation; DR), or dietary restriction plus odour (DR+O).

| *Contrast* | *Estimate* | *SE* | *t-ratio* | *p-value* |
| --- | --- | --- | --- | --- |
| F * F+O | -0.0032 | 0.0014 | -2.232 | 0.122 |
| F * DR | -0.0079 | 0.0014 | -5.643 | <0.0001 |
| F * DR+O | -0.0118 | 0.0014 | -8.413 | <0.0001 |
| F+O * DR | -0.0048 | 0.0014 | -3.388 | 0.006 |
| F+O * DR+O | -0.0087 | 0.0014 | -6.158 | <0.0001 |
| DR * DR+O | -0.0039 | 0.0014 | -2.799 | 0.031 |
| df = 94 for all comparisons | | | | |

**Table S9:** Cox mixed effects model summary for nematode heat shock (37˚C) survival data following mating after treatments of *ad libitum* food (F), *ad libitum* food plus odour (F+O), dietary restriction (24-hour starvation; DR), or dietary restriction plus odour (DR+O). Time points included hours 1-9 of heat shock treatment, and each subsequent time point is a day following heat shock treatment.

|  | *Coefficient* | *Exp(Coefficient)* | *Std. Error* | *z* | *p* |
| --- | --- | --- | --- | --- | --- |
| F+O | 0.0923 | 1.097 | 0.144 | 0.64 | 0.522 |
| DR | -3.181 | 0.042 | 0.211 | -15.10 | <0.0001 |
| DR+O | -3.436 | 0.032 | 0.214 | -16.03 | <0.0001 |
| Random effects: Plate.ID | | Variance: 8.54*10^-5^ | | Std. Deviation: 0.0092 | |

**Table S10:** Zero-inflated negative binomial generalised linear mixed model output for analysis of population growth in semi-natural conditions over 21 days following treatment on Day 1 of adulthood of *ad libitum* food (F), *ad libitum* food plus odour (F+O), dietary restriction (24-hour starvation; DR), or dietary restriction plus odour (DR+O).

| **Model: Population Index ~ Treatment * poly(Day, 2) * Block + (1\|Population/Replicate)** | | | | | |
| --- | --- | --- | --- | --- | --- |
|  | *Estimate* | *Std. Error* | *z-value* | | *p-value* |
| (Intercept) | 3.546 | 0.253 | 14.033 | | <0.0001 |
| F+O | -0.006 | 0.349 | -0.017 | | 0.9861 |
| DR | -0.108 | 0.349 | -0.309 | | 0.7573 |
| DR+O | 0.213 | 0.341 | 0.624 | | 0.5325 |
| Day | 5.118 | 2.329 | 2.198 | | 0.0280 |
| Day^2^ | -26.519 | 2.138 | -12.405 | | <0.0001 |
| Block 2 | -2.092 | 0.352 | -5.950 | | <0.0001 |
| Block 3 | 1.101 | 0.382 | 2.883 | | 0.0039 |
| F+O*Day | 16.290 | 3.365 | 4.841 | | <0.0001 |
| DR*Day | 22.880 | 3.333 | 6.865 | | <0.0001 |
| DR+O*Day | 16.501 | 3.247 | 5.082 | | <0.0001 |
| F+O*Day^2^ | 3.894 | 3.131 | 1.244 | | 0.2137 |
| DR*Day^2^ | -3.877 | 3.086 | -1.256 | | 0.2090 |
| DR+O*Day^2^ | -2.036 | 2.909 | -0.700 | | 0.4841 |
| F+O*Block 2 | 0.254 | 0.490 | 0.519 | | 0.6035 |
| DR*Block 2 | 0.489 | 0.490 | 0.997 | | 0.3187 |
| DR+O*Block 2 | 0.180 | 0.485 | 0.371 | | 0.7108 |
| F+O*Block 3 | 0.049 | 0.534 | 0.091 | | 0.9276 |
| DR*Block 3 | -1.007 | 0.526 | -1.914 | | 0.0556 |
| DR+O*Block 3 | -0.149 | 0.519 | -0.287 | | 0.7743 |
| Day*Block 2 | 13.867 | 4.039 | 3.433 | | 0.0006 |
| Day*Block 3 | -7.355 | 3.330 | -2.209 | | 0.0272 |
| Day^2^*Block 2 | 16.912 | 3.300 | 5.124 | | <0.0001 |
| Day^2^*Block 3 | 19.055 | 3.267 | 5.832 | | <0.0001 |
| F+O*Day*Block 2 | -37.531 | 5.389 | -6.964 | | <0.0001 |
| DR*Day*Block 2 | -30.723 | 5.442 | -5.645 | | <0.0001 |
| DR+O*Day*Block 2 | -21.088 | 5.431 | -3.883 | | 0.0001 |
| F+O*Day^2^*Block 2 | 3.504 | 4.685 | 0.748 | | 0.4545 |
| DR*Day^2^*Block 2 | 5.060 | 4.655 | 1.087 | | 0.2770 |
| DR+O*Day^2^*Block 2 | -10.305 | 4.534 | -2.273 | | 0.0230 |
| F+O*Day*Block 3 | -14.964 | 4.781 | -3.130 | | 0.0017 |
| DR*Day*Block 3 | -22.595 | 4.717 | -4.791 | | <0.0001 |
| DR+O*Day*Block 3 | -20.147 | 4.608 | -4.372 | | <0.0001 |
| F+O*Day^2^*Block 3 | -5.999 | 4.597 | -1.305 | | 0.1919 |
| DR*Day^2^*Block 3 | 7.721 | 4.559 | 1.694 | | 0.0903 |
| R+O*Day^2^*Block 3 | 7.378 | 4.379 | 1.685 | | 0.0920 |
| **Zero-inflation model: ~ Day^2^** | | | | | |
| (Intercept) | -0.197 | 0.510 | -0.386 | | 0.6997 |
| Day | -0.275 | 0.054 | -5.065 | | <0.0001 |
| Block 2 | -0.460 | 0.558 | -0.826 | | 0.4090 |
| Block 3 | -1.205 | 0.440 | -2.741 | | 0.0061 |
| *Random effects:* | | *Variance* | |  | *Std. Dev.* |
| Replicate: Population | | 1.782 * 10^-10^ | | 0.00001 | |
|  | Population | 0.542 | | 0.7445 | |

**Table S11:** Type III Anova output for zero-inflated negative binomial generalised linear mixed model (Table S11) for analysis of population growth in semi-natural conditions over 21 days following treatment on Day 1 of adulthood of *ad libitum* food (F), *ad libitum* food plus odour (F+O), dietary restriction (24-hour starvation; DR), or dietary restriction plus odour (DR+O).

|  | $\boldsymbol{\chi}^{\boldsymbol{2}}$ | **df** | ***p*-value** |
| --- | --- | --- | --- |
| (Intercept) | 4.351 | 1 | 0.0370 |
| Treatment | 10.819 | 3 | 0.0128 |
| Day | 153.965 | 1 | <0.0001 |
| Block | 91.333 | 2 | <0.0001 |
| Day^2^ | 153.756 | 1 | <0.0001 |
| Treatment*Day | 9.273 | 3 | 0.0259 |
| Treatment*Block | 39.736 | 6 | <0.0001 |
| Day*Block | 52.179 | 2 | <0.0001 |
| Treatment* Day^2^ | 6.613 | 3 | 0.0853 |
| Block* Day^2^ | 60.263 | 2 | <0.0001 |
| Treatment*Day*Block | 48.332 | 6 | <0.0001 |
| Treatment* Day^2^*Block | 40.688 | 6 | <0.0001 |

**Supplementary Model Materials:**

Selection against age-specific mortality

Selection against age-specific mortality is expressed by Hamilton (1966) as equal to

$$\begin{aligned} s\left( \mu\left( x \right) \right)=-\int_{y=x}^{\infty} e^{-ry}L\left( y \right)m\left( y \right)dy \#\left( S1 \right) \end{aligned},$$

where $L\left( y \right)$, $m\left( y \right)$ are age-specific rates of cumulative survival and fertility (vital rates), and $r$ is the Malthusian rate of population growth (Fisher 1930). We note two important properties of (*S*1).

1. For all pre-reproductive ages $x<x_{A}$, the integrand in (S1) is equal to zero because $m\left( x \right)=0$.
2. Following from #1 and the Euler-Lotka equation, the integration of (S1) over all adult ages must equal +1.

It therefore follows that any change to values $L\left( y \right)$, $m\left( y \right)$ at any age *y* or to $r$ must cause changes to vital rates at the same or other ages and/or changes in *r*.

To model how DR could affect changes in vital rates and how these might alter patterns of natural selection, we assume that a move from $E_{1}$ to $E_{2}$ decreases age-specific fertility by the same factor $1-k$ regardless of age. It follows that age-specific selection in these two environments is equal to

$$\begin{aligned} s\left( \mu\left( x | E_{1} \right) \right)=-\int_{y}^{\infty} e^{-r\left( E_{1} \right)y}L\left( y | E_{1} \right)m\left( y | E_{1} \right)dy \#\left( S2a \right) \end{aligned}$$

and

$$\begin{aligned} s\left( \mu\left( x | E_{2} \right) \right)=-\int_{y}^{\infty} ke^{-r\left( E_{2} \right)y}L\left( y | E_{2} \right)m\left( y | E_{1} \right)dy \#\left( S2b \right) \end{aligned}.$$

Note that the fertility suppression expressed in $E_{2}$ must either increase cumulative survival or reduce the population growth rate (or both) in order meet the mathematical requirement that the integration equals +1. Biologically, this can be justified by the direct reduction of population growth (which will inflate the value of the exponential term). Furthermore, if there are costs to reproduction that are manifested on mortality at the next time interval, then a relaxation of mortality costs associated with supressed reproduction can increase $L\left( y | E_{2} \right)$ at later ages, and this will inflate the integrand.

We can account for mortality costs of reproduction by allowing for the production of a new offspring to increase mortality at that age by a non-negative value *c*. We model environmental specificity of these costs by subtracting from these a factor *I* in E1 and adding that same value in E2. If, for example, there is a greater survival cost of a reproductive event in the DR environment then with *ad libitum*, then $I>0$. For each age and environment, the total change to the population age-specific mortality rate caused by fertility (the realized costs) is proportional to the corresponding fertility,

$$\begin{aligned} \Delta\mu\left( x | E_{1} \right)=\left( c-I \right) m\left( x | E_{1} \right)\#\left( S3a \right) \end{aligned}$$

$$\begin{aligned} \Delta\mu\left( x | E_{2} \right)=k\left( c+I \right)m\left( x | E_{1} \right)\#\left( S3b \right) \end{aligned}.$$

If we benchmark age-specific mortality in E2 by that in E1, then the former is expressed as in terms of the mortality and fertility of the latter,

$$\begin{aligned} \mu\left( x | E_{2} \right)=\mu\left( x | E_{1} \right)+\delta m\left( x | E_{1} \right) \#\left( S4 \right), \end{aligned}$$

where $\delta=I\left( k+1 \right)-c\left( 1-k \right)$; this is the environmental difference in realized fertility costs.

Strategy for determining when fertility suppression enhances selection for adult lifespan

In principle, one can ask directly how selection changes with age *differently* in the two environments by directly taking the ratio of (S2a,b). We do that for using *C elegans* data derived from our experiment (see Fig X below). However, it is not clearly possible to make general theoretical predictions directly from this ratio owing to the integration functions. Nevertheless, one can make useful predictions using the ratios of the integrands that are specific to common ages. If the ratio of the integrands in (S3b) to (S3a) *increases* with age, then selection against age-specific mortality weakens with age more slowly in $E_{1}$ than in $E_{2}$. If this condition is met, then selection will work more strongly in $E_{2}$ both to resist the evolution of actuarial senescence and to enhance adult lifespan (because selection for mortality at each adult age is stronger in $E_{2}$ than in $E_{1}$). This ratio of integrands is simply

$$\begin{aligned} R\left( y \right)=ke^{\left( r\left( E_{1} \right)-r\left( E_{2} \right) \right)y} \frac{L\left( y | E_{2} \right)}{L\left( y | E_{1} \right)}\#\left( S5 \right). \end{aligned}$$

Because the model does not recognize any feature that might change either the age of reproductive onset or survival to that point, the cumulative lifespan ratio can be restated in terms of adult age-specific mortality. Because $L\left( y \right)=e^{-\int_{0}^{x_{A}} \mu\left( v \right)dv}$, the cumulative survival ratio in (S3) is equal to $\gamma=e^{\int_{x_{A}}^{y} \left( \mu\left( v | E_{2} \right)-\mu\left( v | E_{1} \right) \right)dv}$. We can incorporate costs of reproduction into the ratio of cumulative survival by substituting (S4) into this, yielding

$$\begin{aligned} \gamma=e^{\delta\int_{x_{A}}^{y} m\left( v | E_{1} \right)dv} \#\left( S6 \right) \end{aligned}$$

Following the logic described above, we can infer that selection for adult lifespan is always stronger in $E_{2}$ than in $E_{1}$ if the ratio given in (S5) has a non-decreasing monotonic relationship with *y*. The conditions sufficient to satisfy this requirement correspond to those that find the first derivative of $R\left( y \right)$ taken with respect to *y* to be always greater than zero.

Deriving the conditions that enhance selection for adult lifespan

Following this logic for all values of *y*, the critical values are given by the inequality

$$\begin{aligned} ke^{\left( r\left( E_{1} \right)-r\left( E_{2} \right) \right)y} \left( \frac{d\gamma}{dy}+\left( r\left( E_{1} \right)-r\left( E_{2} \right) \right)\gamma\right)>0 \#\left( S7 \right) \end{aligned},$$

where $\frac{d\gamma}{dy}=\gamma\delta m\left( y | E_{1} \right)$. Substituting into (S7), this is

$$\begin{aligned} k\gamma e^{\left( r\left( E_{1} \right)-r\left( E_{2} \right) \right)y} \left( \delta m\left( y | E_{1} \right)+r\left( E_{1} \right)-r\left( E_{2} \right) \right)>0 \#\left( S8 \right) \end{aligned},$$

and because $k,\gamma$ and $e^{x}$must be greater than zero for all *x*, this inequality simplifies to

$$\begin{aligned} \delta m\left( y | E_{1} \right)+r\left( E_{1} \right)-r\left( E_{2} \right)>0 \#\left( S9 \right) \end{aligned}.$$

This difference in population growth rates can be expressed in terms of average birth and death rates taken over all individuals in each environment. In general, $r=b-d$, and we can characterize both growth rates in terms of birth and death rates in $E_{1}$ following relationships established before. Owing to the effects of fertility suppression, $b\left( E_{2} \right)=kb\left( E_{1} \right)$. Furthermore, $\mu\left( E_{2} \right)=\mu\left( E_{1} \right)+\delta b\left( E_{1} \right)$. Taking the exponential of both sides of the latter expression and recognising that mortality rates are the natural logarithms of death rates, it follows that $d\left( E_{2} \right)=d\left( E_{1} \right)e^{\delta b\left( E_{1} \right)}$. Substituting these relationships into the difference in population growth rates yields

$$\begin{aligned} r\left( E_{1} \right)-r\left( E_{2} \right)= b\left( E_{1} \right)\left( 1-k \right)-d\left( E_{1} \right)\left( 1-e^{\delta b\left( E_{1} \right)} \right) \#\left( S10 \right) \end{aligned}.$$

Substituting (S10) in (S9) defines the necessary and sufficient conditions for selection for adult lifespan to be stronger in $E_{2}$ than in $E_{1}$ expressed in terms of birth and death rates,

$$\begin{aligned} \delta m\left( y | E_{1} \right)+b\left( E_{1} \right)\left( 1-k \right)-d\left( E_{1} \right)\left( 1-e^{\delta b\left( E_{1} \right)} \right)>0 \#\left( S11 \right). \end{aligned}$$

Solving for *k* finds its critical value

$$\begin{aligned} k<1+\frac{\delta m\left( y | E_{1} \right)-d\left( E_{1} \right)\left( 1-e^{\delta b\left( E_{1} \right)} \right)}{b\left( E_{1} \right)} \#\left( S12 \right). \end{aligned}$$

We can evaluate the conditions defined in (S11-12) to examine some simple cases.

Case 1: No environmental differences in fertility costs, $\delta=0$.

This can be because there are no fertility costs in either environment $\left( c,I=0 \right)$ or, more generally, the realized fertility costs are the same in both environments, $k\left( c+I \right)=c-I$. Here (S12) simplifies to $k<1$. Because this condition is one of our premises ($E_{2}$ suppresses fertility), one can conclude that selection is always enhanced in this simple case. This happens because the resulting reduction in population growth rate shifts strengthens late-life selection (compared to selection at the same ages in the high fertility population).

Case 2: There is a greater realized mortality cost to reproduction in $E_{2}$, $\delta>0$.

Eq (S11) demonstrates that selection is stronger in $E_{2}$ if the sum of three terms is positive. If $\delta>0$ (our premise), then the first term is positive. The second term is always positive. The third term can be re-arranged as $+d\left( E_{1} \right)\left( e^{\delta b\left( E_{1} \right)}-1 \right)$; this also must be positive if $\delta>0$, which again is our premise. Positive differential costs to reproduction can happen when the per-reproduction costs to fertility are much higher in the population with suppressed fertility. Specifically, $\delta>0$ if and only if $I>\frac{\left( 1-k \right)}{\left( 1+k \right)}c$.

Eq (S11) is too complex to provide a more insightful general description of threshold conditions that define all cases where selection strengthens in $E_{2}$. However, we can contrive a simple case that demonstrates that conditions can be satisfied when there are *less* realized mortality costs to reproduction in $E_{2}$ ($\delta<0$).

Case 3: Fertility is age-independent

In this case, we can set $m\left( y | E_{1} \right)=b\left( E_{1} \right)$ and substitute into (S11). Re-arranging, this yields this condition,

$$\begin{aligned} \delta>\frac{ln\left( \left( k-\delta\right)b\left( E_{1} \right)-b\left( E_{1} \right)+d\left( E_{1} \right) \right)-ln\left( d\left( E_{1} \right) \right)}{b\left( E_{1} \right)} \#\left( S13 \right). \end{aligned}$$

If the right-hand side of (S13) can be shown to be less than zero and $\delta$, then this implies that negative values of $\delta$ exist that satisfy inequality (S13). This new inequality to be tested simplifies to $k-\delta<1$, and this means that selection is strengthened in $E_{2}$ when

$$\begin{aligned} 0>\delta>-\left( 1-k \right)\#\left( S14 \right), \end{aligned}$$

where $1-k$ is the fractional suppression of fertility. These conditions can be characterized in terms of mortality costs per reproduction by substituting $\delta=I\left( k+1 \right)-c\left( 1-k \right)$, and this yields the following conditions.

$$\begin{aligned} 0>I\left( k+1 \right)-c\left( 1-k \right)>-\left( 1-k \right) \#\left( S15 \right). \end{aligned}$$

All that remains is to show by example that a solution exists. Parameters $k,I=0.5$ and $c=2$ satisfy these inequalities $\left( i.e., 0>-0.25>-0.5 \right)$.
